# Age-related changes in social behaviours in the 5xFAD mouse model of Alzheimer’s disease

**DOI:** 10.1101/471714

**Authors:** Filip Kosel, Paula Torres Munoz, J. Renee Yang, Aimee A. Wong, Tamara B. Franklin

## Abstract

In addition to memory impairments, patients with Alzheimer’s disease (AD) exhibit a number of behavioural and psychological symptoms that can affect social interactions over the course of the disease. While altered social interactions have been demonstrated in a number of mouse models of AD, many models only recapitulate the initial stages of the disease, and these behavioural changes have yet to be examined over the course of disease progression. By performing a longitudinal study using the 5xFAD mouse model, we have demonstrated that transgenic females exhibit progressive alterations in social investigation compared to wild-type controls. Transgenic females exhibited an age-related reduction in interest for social odours, as well as reduced investigative behaviours towards novel conspecifics in a novel environment. However, transgenic mice exhibited no obvious olfactory deficits, nor any changes in scent-marking behaviour compared to wild-type controls, indicating that changes in investigative behaviour were due to motivation to engage with a social stimulus. This evidence suggests that transgenic 5xFAD females exhibit increased social anxiety in novel environments compared to wildtype controls. Overall, transgenic 5xFAD female mice mimic some features of social withdrawal observed in human AD patients suggesting this strain may be suitable for modelling aspects of the social dysfunction observed in human patients.

## 1. Introduction

In addition to the memory deficits and cognitive impairments commonly associated with Alzheimer’s disease (AD), the progressive neurodegeneration also results in a number of behavioural and psychological symptoms of dementia (BPSD); these include delusions, hallucinations, agitation/aggression, anxiety, elation/euphoria, apathy/indifference, disinhibition, irritability, and aberrant motor behaviour [1,2]. Approximately 80% of patients exhibit at least one BPSD following the onset of cognitive symptoms, with agitation/aggression, irritability, depression, anxiety, and apathy being the most common [3,4]. BPSD can have a significant impact on social interactions, with social withdrawal being highly prevalent in AD patients, even prior to diagnosis [3-9]. Behavioural disturbances, including agitation/aggression, are a leading factor in institutionalization, and placement in a care facility can affect AD patients by reducing meaningful social interactions and leading to potential infantilization by care staff [10-15].

Given the importance of social interactions on the well-being of AD patients, a number of genetic mouse models have been used to examine effects of AD-related pathology on social behaviours. Results have indicated impaired social recognition memory in 21-month old transgenic Tg2576 females and increased aggression in a resident-intruder task in 7-month old transgenic Tg2576 males [16-18]. Evidence for increased sociability in the 3xTg-AD mouse model has also been presented [19], but the Tg2576 strain exhibits no effect of AD pathology on sociability [17]. Overall, these results indicate that some mouse models of AD exhibit changes in social behaviour that include altered aggression and sociability. However, many transgenic mouse models of AD (e.g., 3xTg-AD, APP/PS1, and Tg2576) exhibit a slow rate of amyloid beta (Aβ) accumulation and can only model the initial phase of the disease, which does not include widespread neurodegeneration [20].

By comparison, the 5xFAD strain exhibits rapid development and progression of amyloid plaques and neurodegeneration, allowing effects to be tracked across early-, mid-, and late-stage disease progression [20,21]. There is increasing evidence that 5xFAD mice exhibit a range of social deficits that, in some cases, mirror the social symptoms presented by AD patients. Transgenic 5xFAD 9-month-old male mice and 12-month-old male and female mice demonstrate deficits in nest building, a behaviour that has been associated with affiliative behaviours [22,23]; however, they also engage in more homecage social behaviours than wild-type controls [24]. Transgenic 5xFAD mice also exhibit impaired social recognition memory at 9-months of age [24], consistent with memory deficits exhibited in this strain and human AD patients.

While these results suggest that the 5xFAD strain exhibit social deficits, a comprehensive investigation of social behaviours in this strain has not yet been performed. The present study aimed to characterize a range of social behaviours in the 5xFAD strain in a genotype- and age-dependent manner. Specifically, this study examined responses to social odour cues, sociability and social novelty preference, and free social interaction. Overall, it was hypothesized that transgenic 5xFAD mice would exhibit abnormal responses to social odour cues and altered sociability, social novelty preference, and free social interaction compared to wild-type controls in an age-dependent manner.

## 2. Material and Methods

### 2.1 Animals

Mice were the progeny of male hemizygous C57BL/6J x SJL/J FN 5xFAD (B6SJL-Tg (APPSwFILon, PSEN1*M146L*L286V) 6799Vas/Mmjax) and female WT C57BL/ 6J x SJL/J F1 mice (B6SJLF1/J, JAX stock # 100012) obtained from Jackson Laboratories (Bar Harbor, ME, USA) and bred in our laboratory.

All mice were housed in a colony room on a 12:12 reversed light:dark cycle, with lights off from 09:30 – 21:30. Mice were housed in standard polysulfone cages (30 x 19 x 13 cm; model PC7115HT; Allentown Caging Inc., Allentown, NJ, USA) containing wood-chip bedding (FreshBed; Shaw Resources, Shubenacadie, NS, Canada), a metal cage top containing *ad libitum* food (Laboratory Rodent Diet #5001; Purina LabDiet, St. Louis, MO, USA) and filtered tap water, with two black, opaque, polymer enrichment tubes (4 cm diameter, approx. 8 cm long). Cages were topped with a micro-isolator filter to reduce the spread of airborne contaminants and diseases. Cages were changed weekly, just prior to the end of the light cycle. To reduce instability of dominance hierarchies, a small amount of bedding was transferred from the old cage to the new cage.

Mice were genotyped for the PSEN transgene and the phosphodiesterase-6b retinal degeneration-1 (Pde6b^rd1^) allele by standard PCR procedures (Jackson Laboratories, Bar Harbor, ME, USA), and the primers listed in Table 1, using tissue samples obtained from ear punches of mice between P14 and P18. Mice were weaned between P23 and P26, and were group-housed in same-sex, same-genotype, same-litter cages of 2-4 animals. Female transgenic and wild-type mice were tested at 13 weeks (3 months), 26 weeks (6 months), 39 weeks (9 months), and 52 weeks (12 months) of age. Mice exhibiting stereotypy (continuously running in circles, doing backflips, jumping in one corner of the cage) were excluded from testing to ensure that these behaviours did not affect results. Mice that were homozygous recessive for Pde6b^rd1^, which is common in this mouse strain and leads to retinal degeneration, were not used to ensure that visual impairments did not affect results. Singly-housed subjects were also excluded from testing. See Table 2 for sample sizes at each age. All experimental procedures were performed in accordance with the guidelines published by the Canadian Council on Animal Care and were approved by Dalhousie University Committee on Laboratory Animals (Protocol #16-021).

**Table 1.**
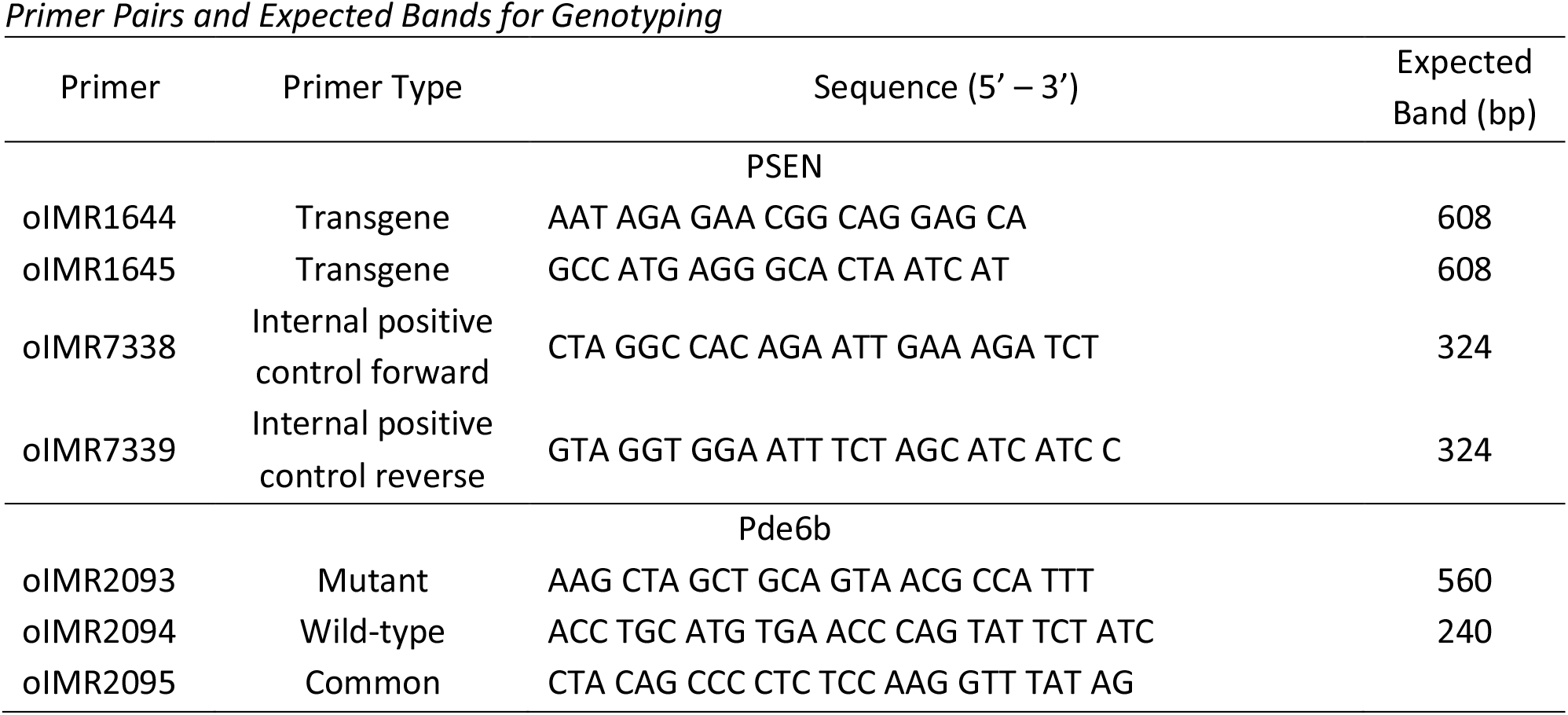
Primer Pairs and Expected Bands for Genotyping

**Table 2.** Sample Sizes by Age and Genotype

| Genotype | 3 Months | 6 Months | 9 Months | 12 Months |
| --- | --- | --- | --- | --- |
| Transgenic | 24 | 22 | 21 | 20 |
| Wild-Type | 17 | 17 | 17 | 14 |

### 2.2 Behavioural Procedures

#### 2.2.1 Testing Schedule and Conditions

All testing was performed within an 11-hour window, starting a minimum of 30 minutes after onset of the dark phase, and ending at least 30 minutes prior to onset of the light phase. Tasks were performed as follows: Day 1, olfactory habituation/dishabituation; Day 3, urine marking; Day 5, sociability and social novelty preference; Day 8, free social interaction. Testing was performed in a quiet room under dim light conditions. All behavioural tasks were recorded using the Biobserve Viewer 3.0 program (Biobserve GmbH, Bonn, Germany).

#### 2.2.2 Olfactory Habituation/Dishabituation

The apparatus consisted of a standard polysulfone cage (30 x 19 x 13 cm; identical to the home cage) containing wood chip bedding and topped with a metal cage top. Wood-handled cotton swabs (15 cm long; Puritan Medical Products Company, Guildford, MN, USA) were inserted between the bars of the cage top and held in place with a binder clip, with the swab and approximately 3 cm of the handle protruding into the cage approximately 5 cm above the base of the cage. Cotton swabs were inserted from the handle end to avoid transferring scent from the swab to the metal cage top.

Stimuli consisted of the following scents, in order of presentation: *1)* distilled water (neutral control); *2)* imitation almond extract (1:100 dilution; La Cie McCormick Canada, London, ON, Canada); *3)* imitation banana extract (1:100 dilution; La Cie McCormick Canada); *4)* unfamiliar same-sex cage 1; *5)* unfamiliar same-sex cage 2. All stimuli were prepared on the day of testing and were only used for that day. Dilutions were performed using the same water as the neutral control, and stimuli were prepared on the day of testing. Non-social odour cues were made by preparing the dilution in a 15 mL conical centrifuge tube, and briefly immersing the swab in the dilution prior to use. Each of the non-social odour cues used throughout the experiment came from the same original bottles, which were stored at 4°C and vortexed prior to use. Social odour cues were prepared by swabbing the bottom of an unfamiliar, same-sex cage that had not been changed for at least three days prior and each stimulus cage contained at least two animals. Any bedding material remaining on the swabs was brushed off, and swabs were placed in a zipper storage bag until use.

The procedure was performed as described by Yang and Crawley [25]. Subjects were placed into a clean cage, the metal cage top containing a clean, dry cotton swab was placed on top, and the cage was covered with a microisolator top; subjects were allowed to habituate to this apparatus for 30 minutes prior to testing to reduce the novelty of the cotton swab. During this time, subjects remained in the colony room. All subjects were removed from home-cages and placed into testing cages at the same time and were returned to the home-cage once all cage-mates had completed testing.

Following the 30-minute habituation period, the dry swab was removed, and subjects were moved to the testing room. A swab was prepared with water alone and was placed into a metal cage top; this top was placed onto the cage, and the recording was started. Each recording period was 2 minutes long, and during this time the experimenter left the room and prepared a second cage top with the next stimulus. When the recording finished, the cage top containing the previous stimulus was replaced with the next stimulus, and the next recording was started. This process was repeated until all 15 stimuli had been presented.

Following testing, subjects were returned to the colony room. Subjects remained in the testing cage until all cage-mates had been tested before being returned to their home-cage. Subjects not undergoing habituation or testing were given a metal cage top containing a bottle of water to prevent dehydration.

Videos were scored manually for time in seconds spent sniffing during each presentation of each odour, as described by Yang and Crawley [25]. Sniffing was defined as orientation of the snout towards the cotton swab, with the swab 2 cm or less from the snout. Raters were blind to the genotypes of the animals.

Data analysis was performed in R version 3.4.1 (R Foundation for Statistical Computing, Vienna, Austria) using 95% confidence intervals of Cohen’s d for genotype and habituation/dishabituation effects. Outliers were not excluded.

#### 2.2.3 Urine Marking

The apparatus consisted of a polycarbonate arena, 40 x 40 x 60 cm, with four opaque white walls and a colourless and transparent floor. The bottom of the arena was lined with a sheet of clean, absorbent paper (Strathmore 400 Series Recycled Drawing Paper; Strathmore Artist Papers, Neenah, WI, USA), similar to the apparatus described by Roullet, Wohr, and Crawley [26]. Odour stimuli were prepared by placing 40 μL spots of the odour cue on the paper in each corner of the arena for a total of 4 spots (160 μL), similar to that described by Ninomiya and Kimura [27]. Each sheet contained only a single odour cue placed in each corner, with no odours present during habituation.

Stimuli consisted of three odour cues: *1)* imitation vanilla extract (1:100 dilution; La Cie McCormick Canada); *2)* same-sex conspecific urine; *3)* opposite-sex conspecific urine. Vanilla scent (non-social cue) was prepared on the day of testing using distilled water. The vanilla extract used throughout the experiment came from the same original bottle and was stored at 4°C and vortexed prior to use. Urine was collected from unfamiliar, sexually-naive, adult, wild-type C57B6SJLF2 mice. Urine collection was performed similarly to that described by Yang and Crawley [25]: mice were scruffed, a clean 1.5 mL microcentrifuge tube was held near the genitals, and the abdomen was palpated gently to promote urination. Urine was collected from at least four mice over the course of seven days. Following collection, urine was refrigerated at 4° C, then pooled and divided into 1 mL aliquots. Pooling was done to ensure odour cues are consistent across testing, and do not vary across stage of estrous or dominance of sample provider. Aliquots were then frozen at −80° C until use. Odour stimuli were kept on ice during testing.

For testing, subjects were transferred to a clean holding cage containing bedding and a food pellet, and a micro-isolator cover was placed on top. All subjects were removed from home-cages and placed into holding cages at the same time. They were not returned to the home-cage until all cage-mates had completed testing.

An arena was prepared with a clean sheet of paper with no odour, and the subject was placed into the arena and allowed to freely explore for 5 minutes to habituate. During this time, a second arena was prepared with the next paper sheet and stimulus; order of presentation of each stimulus was counterbalanced across subjects and across ages, and arenas were prepared in a separate room from testing to minimize odour in the testing room. At the end of the 5-minute habituation period, the subject was returned to its holding cage, the arena with the habituation sheet was removed, and the arena containing the next stimulus was put in its place. The mouse was then placed in the next arena and allowed to freely explore for 15 minutes. The first arena—containing the habituation sheet—was then moved to a separate room away from the testing area; this was done to reduce odours from reaching the testing room. A third arena was then prepared with the next stimulus, and the process was repeated until the mouse had performed one 5-minute habituation trial and three 15-minute testing trials. Sheets were allowed to dry for 15 minutes immediately following testing, and then were carefully removed from the apparatus and placed on a flat surface to further dry for approximately 24 hours. In between trials, arenas were cleaned with Fisherbrand Sparkleen 1 (Fisher Scientific, Pittsburgh, PA, USA) and room-temperature water, then thoroughly dried with paper towels.

Following testing, subjects were returned to the colony room. Subjects remained in the holding cage until all cage-mates had been tested before being returned to their home-cage. Subjects not undergoing testing were given a metal cage top containing a bottle of water to prevent dehydration.

Scoring was done by examining the surface area of subject urine covering the sheets. After drying, sheets were sliced into quarters (approximately 20 x 20 cm), then were imaged using a Bio-Rad ChemiDoc MP Imaging System and Image Lab Version 5.1 (Bio-Rad Laboratories, Hercules, CA, USA) using the following settings: blue light illumination; 530/28 filter; 50 msec exposure time. These settings were previously determined to provide the best fluorescence of urine marking while minimizing background fluorescence. All images were taken at a uniform zoom level to allow comparison of images at varying time points. Images were then exported as TIFF files for analysis in ImageJ Version 1.51 (U.S. National Institutes of Health, Bethesda, MD, USA). Image thresholding was used to identify fluorescence, using the following settings: 14500 lower limit; 65535 upper limit; MaxEntropy protocol; B&W output; dark background. As with imaging, these settings were previously found to provide the best differentiation between fluorescence of urine spots and background fluorescence. Following thresholding, a 1 x 1 cm grid was overlaid on the image, and the number of grid squares containing urine spots was counted [26,28-30]. Previous studies [28,29] defined urine pools as any spots encompassing more than 4 square grids (2 x 2 cm); however, placement of the grid may affect whether a spot is marked as a urine pool, and urine pools may shift during handling of the apparatus, thereby affecting their shape. As a result, the present study deemed urine pools to be any spots greater than 4 cm^2^, regardless of shape. Urine pools and spots corresponding to the original locations of the odour stimuli were removed from analysis, and the number of grid squares covered by remaining marks was summed for each sheet.

Data analysis was performed in R version 3.4.1 (R Foundation for Statistical Computing) using multilevel modelling. Subjects that were separated or exhibiting stereotypy were not testing, resulting in missing data at certain ages. In order to make use of the remaining data, multilevel modelling was chosen for data analysis over repeated-measures ANOVA (which requires that all subjects have data for each time-point). Genotype was used as the between-subjects factor, while age and stimulus were used as within-subjects factors. The effect of subject was used as the error term. A preliminary examination indicated that the data exhibited a negative binomial distribution; as a result, a logarithm transformation was used to normalize the residuals in order to perform accurate data analysis. Genotype effects were examined using 95% confidence intervals of Cohen’s d.

Due to the presence of liquid on the sheet (including odour stimuli and urine pools from the subjects), subjects could artificially increase the surface area of the fluorescence by dragging their tails through or stepping in liquid. To avoid bias by the rater and prevent removal of actual urine marking, these marks were not excluded from analysis; instead, outliers (sheets where the number of grid squares covered was over two standard deviations from the mean) were excluded from analysis. Outliers were determined by calculating the mean and standard deviation for each genotype, age, and stimulus separately, then comparing the results of each sheet to those values.

#### 2.2.4 Sociability and Social Novelty Preference

The apparatus was a three-chamber box similar to that described by Moy *et al* [31]. The apparatus consisted of a clear, acrylic box, 69 x 20 x 20 cm, divided into three compartments of equal size (23 x 20 x 20 cm) by two clear, acrylic walls. Access to each compartment was through a 6 x 6 cm, floor-level opening in the middle of the dividing walls. Opaque sections of acrylic were used as barriers between the three chambers. The floor of the apparatus was lined with the same wood-chip bedding as used in the home-cages, and a round wire cage (Galaxy Cup; Spectrum Diversified Designs Inc., Streetsboro, OH, USA) was used in each of the two outer chambers to contain stimuli. White, opaque, 500 mL HDPE bottles (Nalge Nunc International Corporation, Rochester, NY, USA) filled with water were placed on top of the Galaxy Cups to prevent subjects from climbing on top.

Prior to testing, subjects were transferred to a clean holding cage containing wood-chip bedding and a food pellet, and a micro-isolator cover was placed on top. All subjects were removed from home-cages and placed into holding cages at the same time. They were not returned to the home-cage until all cagemates had completed testing. Stimuli mice were also transferred to holding cages for testing.

The sociability and social novelty tasks were performed similarly to that described by Gunn *et al* [32]. Each trial began with the opaque barriers in place, blocking access between the three chambers. Stimuli were placed under the Galaxy Cups, with water bottles on top, and the subject was placed in the centre chamber. Once the subject and stimuli were in place, both barriers were removed simultaneously, and the subject was allowed to freely explore the apparatus for the duration of the trial; habituation trials were 5 min in length, while testing trials were 10 min in length. At the end of the trial, the subject was coaxed into the centre chamber and both barriers were replaced. To reduce stress due to repeated handling, subjects were allowed to remain in the centre chamber of the apparatus while the stimuli were quickly replaced.

Habituation trials included no stimulus under either Galaxy Cup. The first testing trial began one minute after the habituation trial, and the stimuli consisted of a small, unfamiliar plastic animal toy (non-social stimulus) placed under one Galaxy Cup, and an unfamiliar, same-sex, wild-type stimulus mouse placed under the Galaxy Cup in the opposite chamber; this trial was used to examine sociability, and the side of presentation of the toy and mouse were counterbalanced across subjects and across ages. All stimulus mice were habituated to the Galaxy Cups for 15 minutes prior to testing. The second testing trial began one minute after the first testing trial. For the second testing trial, the toy was removed, and a second unfamiliar, same-sex, wild-type stimulus mouse was placed in its location; the first stimulus mouse remained, and due to being exposed to the subject during the previous trial, was now considered the familiar mouse. This trial examined social novelty preference.

Following testing, subjects were returned to the colony room. Subjects remained in the holding cage until all cage-mates had been tested before being returned to their home-cage. Subjects not undergoing testing were given a metal cage top containing a bottle of water to prevent dehydration.

The apparatus was cleaned in between subjects, with bedding removed and any remaining particles vacuumed out. The apparatus was cleaned using Fisherbrand Sparkleen 1 (Fisher Scientific) and roomtemperature water, then thoroughly dried with paper towels. Between testing different cohorts of subjects, Galaxy Cups were cleaned with Fisherbrand Sparkleen 1 and room-temperature water then dried with paper towels, to reduce any possible odours from urine marking.

Videos were manually scored for duration (in seconds) spent in each chamber during each trial, as well as duration (in seconds) of interaction with each stimulus. Subjects were considered to have entered a chamber as soon as their head and forelimbs were across the threshold into the chamber. Interaction with a stimulus was defined as orientation of the snout towards the Galaxy Cup, with the snout 1 cm or less away from the Galaxy Cup. Interaction duration was used to calculate preference ratios for the novel stimulus animal over the novel toy (sociability), or the novel stimulus animal over the familiar stimulus animal (social novelty preference and social recognition memory). Preference ratios were calculated by dividing the time spent interacting with the novel stimulus animal over the total time spent interacting with either stimulus. Additionally, to ensure that preference ratio differences were not due to overall differences in exploration, the total time (in seconds) spent interacting with both stimuli was also calculated. Previous work by Fairless *et al* [33] indicated that cylinder time (time spent investigating the cylinder/cage containing a stimulus) was a more consistent measure of sociability than time spent in chamber; as a result, all analyses were performed using investigation time.

Data analysis was performed in R version 3.4.1 (R Foundation for Statistical Computing) using multilevel modelling. Data was analyzed for preference ratio and investigation time of both stimuli for the entire trial. Due to expected age-related effects, 12-month data was also examined for preference ratio and total investigation time in 2-minute intervals (bins), as well as latency to first approach to both stimuli. Genotype was used as the between-subjects factor, and age was used as the within-subjects factor. The effect of subject was used as the error term. Trials were analyzed independently: Trial 1 examined sociability (social vs non-social stimuli) and Trial 2 examined social novelty preference (novel vs familiar social stimuli). Genotype effects were examined using 95% confidence intervals of Cohen’s d. Outliers were not removed from analysis.

#### 2.2.5 Free Social Interaction

The apparatus consisted of a 38 x 38 x 40 cm box, with the floor raised 5 cm off of the base. Three of the walls were made of plywood and painted beige, and the fourth wall was made of clear acrylic to allow for video recording from the front. The floor was made of clear acrylic to facilitate cleaning between animals; a sheet of black paper was placed immediately below the acrylic so that it appeared solid.

Prior to testing, subjects were transferred to a clean holding cage containing wood-chip bedding, a food pellet, and a micro-isolator cover was placed on top. All subjects were removed from home-cages and placed into holding cages at the same time, and were not returned to the home-cage until all cagemates had completed testing. Stimulus mice were also transferred to holding cages for testing.

The protocol was largely based on existing free social interaction protocols [19,33-35]. The subject was placed into the apparatus and allowed to habituate by freely exploring for five minutes. Following habituation, subjects remained in the apparatus while an unfamiliar, same-sex, wild-type stimulus mouse was placed into the apparatus; the mice were then allowed to freely interact for 10 minutes.

Following testing, subjects were returned to the colony room. Subjects remained in the holding cage until all cage-mates had been tested before being returned to their home-cage. Subjects not undergoing testing were given a metal cage top containing a bottle of water to prevent dehydration. The apparatus was cleaned between subjects using Fisherbrand Sparkleen 1 (Fisher Scientific) and room-temperature water, then thoroughly dried with paper towels.

Videos were manually scored for duration (in seconds) of investigative (follow; orofacial sniffing; anogenital sniffing) and self-grooming behaviours. Affiliative (sitting in contact; allogrooming) and aggressive (chase; mounting; attack) behaviours were also tracked, but were not exhibited by subjects and were removed from analysis. Definitions of these behaviours were adapted from Arakawa, Blanchard, & Blanchard [36] and Grant and Mackintosh [37], and were as follows: follow (slow speed locomotion towards the other animal that is also moving); orofacial sniffing (orientation of the snout towards and nearly touching the snout of the other animal); anogenital sniffing (orientation of the snout towards and nearly touching the anogenital region of the other animal); self-grooming (all licks given by an animal to itself). Although Grant and Mackintosh [37] classified anogenital sniffing under mating behaviour, it is also used to determine dominance in mice, and as a result it was considered an investigative behaviour in the present study.

Data analysis was performed in R version 3.4.1 (R Foundation for Statistical Computing) using multilevel modelling; see Section 2.2.2.4.4 for rationale. Data was analyzed for total behaviour duration for the entire trial. Due to expected age-related effects, 12-month data was also examined for duration of behaviours in 2-minute intervals (bins). Genotype was used as the between-subjects factor, while age was used as the within-subjects factor. The effect of subject was used as the error term. Due to behaviours being relatively independent of each other, each behaviour was analyzed separately. Genotype effects were examined using 95% confidence intervals of Cohen’s d.

### 2.3 Data and Code Availability

All behavioural videos and processed data are publicly available at the Federated Research Data Repository (https://dx.doi.org/10.20383/101.0126). The raw data and the code used to analyze the data is available from the corresponding author upon reasonable request.

## 3. Results

### 3.1 Olfactory Habituation/Dishabituation

Transgenic 5xFAD females and wild-type controls exhibit olfactory habituation (decrease in sniffing duration) over successive presentations of a given odour cue, as well as olfactory dishabituation (increase in sniffing duration) in response to the first presentation of a different odour cue (Figures 1 and S1).

**Figure 1.**
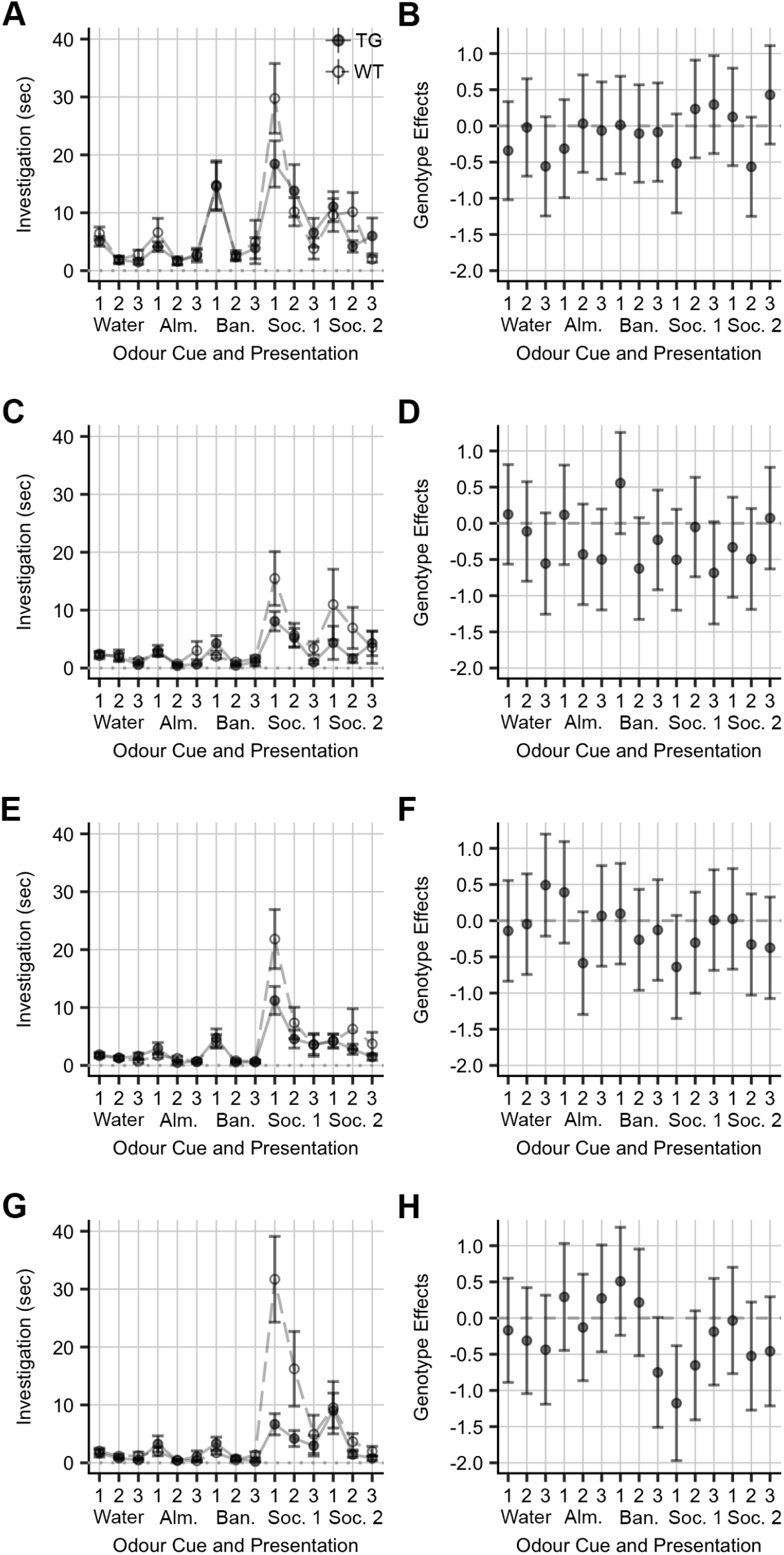
Odour Investigation in the Olfactory Habituation/Dishabituation Task. Mean and standard error for sniffing durations (in sec) for each odour and presentation during the Olfactory Habituation/Dishabituation task for females at **A)** 3 months, **C)** 6 months, **E)** 9 months, and **G)** 12 months of age, as well as 95% CIs for Cohen’s d for genotype effects at **B)** 3 months, **D)** 6 months, **F)** 9 months, and **H)** 12 months of age. Points indicate means, error bars indicate standard error (panels A, C, E, and G) and 95% CIs (panels B, D, F, and H). Effects given for transgenic females relative to wild-type controls. TG = transgenic, WT = wild-type, Alm. = almond cue, Ban. = banana cue, Soc. 1 = first social odour cue, Soc. 2 = second social odour cue.

Compared to wild-type control females, transgenic 5xFAD females typically exhibit an age-related decrease in sniffing durations in response to social odours, particularly for dishabituation to the first social odour (Cohen’s d 95% CIs [−1.20, 0.17], [−1.20, 0.19], [−1.35, 0.07], and [−1.97, −0.38] for genotype effects on sniffing durations for social odour cue 1 at 3, 6, 9, and 12 months, respectively; Table 3 and Figure 1).

**Table 3.**
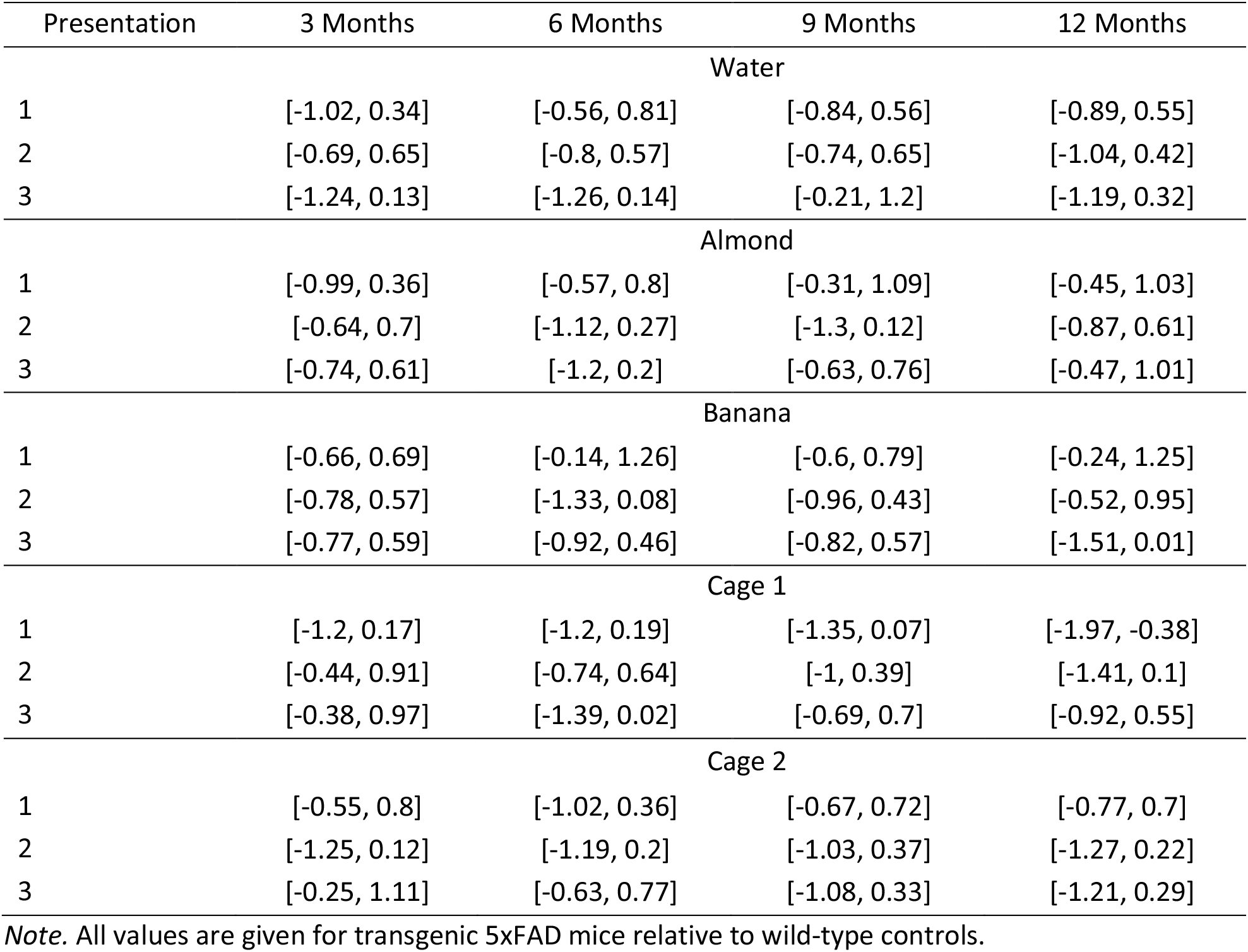
95% Confidence Intervals for Cohen’s d for Genotype Effects on Sniffing Duration for Each Odour Cue During the Olfactory Habituation/Dishabituation Task from 3 to 12 Months of Age.

### 3.2 Urine Marking

Analysis of females from 3 to 12 months of age indicated a main effect of age (χ^2^ (1) = 45.20, *p* < .001) and odour cue (χ^2^ (2) = 83.73, *p* < .001), with increased urine marking in response to social cues over non-social cues (with no difference in urine marking to same- vs opposite-sex urine), and an overall decrease in urine marking with age, particularly from 6 to 9 months of age (Figure 2 and Table 4). There was no effect of genotype or genotype by age interaction (Figure 2 and Table 4).

**Figure 2.**
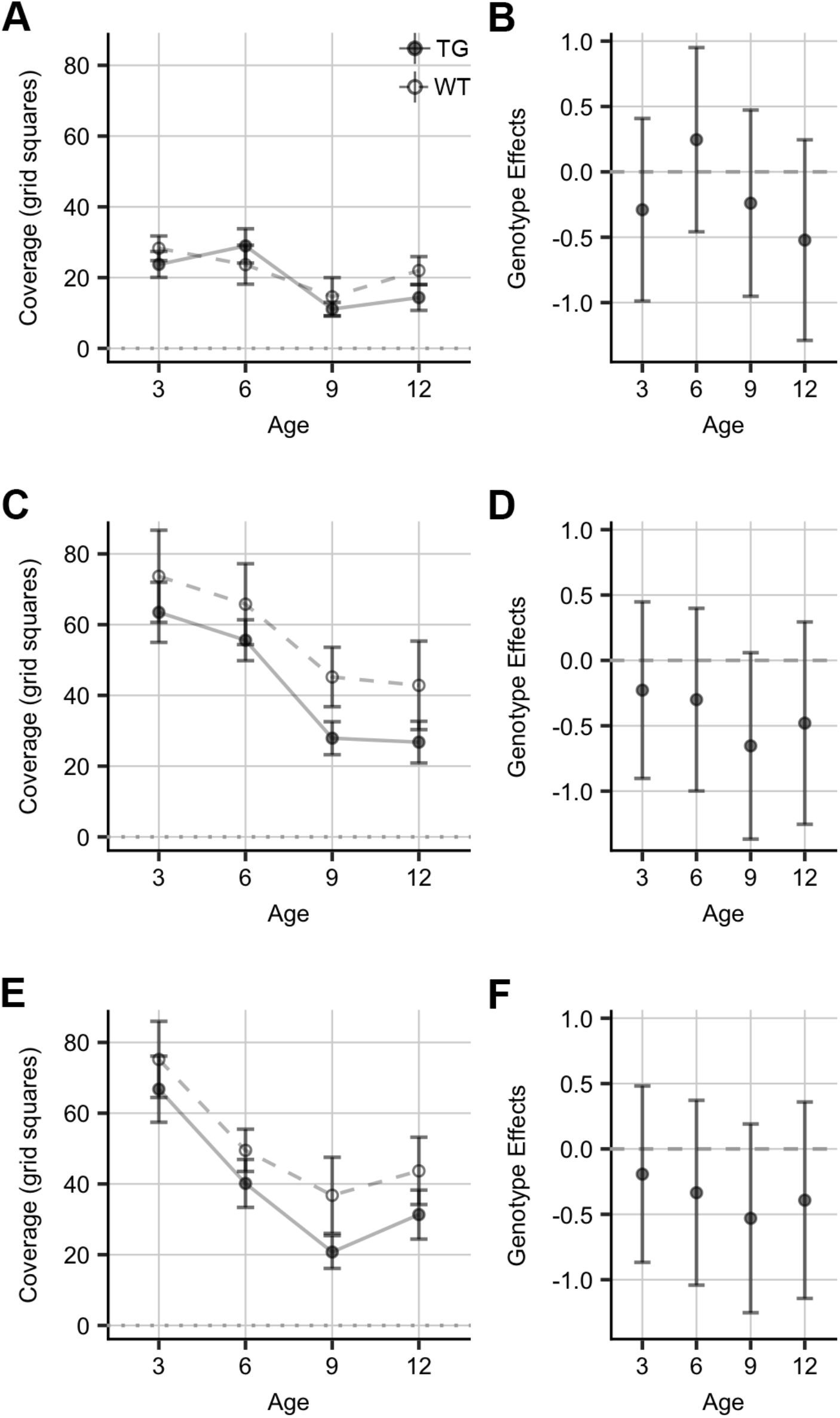
Scent Marking Behaviour in the Urine Marking Task. Mean and standard error for coverage (in grid squares) for **A)** neutral, **C)** same-sex urine, and **E)** opposite-sex urine cues, as well as 95% CIs for Cohen’s d for genotype effects for **B)** neutral, **D)** same-sex urine, and **F)** opposite-sex urine cues, during the Urine Marking task for females from 3 to 12 months of age. Points indicate means, error bars indicate standard error (panels A, C, and E) and 95% CIs (panels B, D, and F). Effects given for transgenic females relative to wild-type controls. TG = transgenic, WT = wild-type.

**Table 4.** Analysis of Deviance for the Urine Marking Task from 3 to 12 Months of Age

| Effect | $\chi^2$ | df | *p* |
| --- | --- | --- | --- |
| Genotype | 3.38 | 1 | .066 |
| Age | 45.20 | 1 | **.000** |
| Odour Cue | 83.73 | 2 | **.000** |
| Genotype:Age | 0.97 | 1 | .324 |
| Genotype:Odour Cue | 0.92 | 2 | .631 |
| Age:Odour Cue | 1.03 | 2 | .598 |
| Genotype:Age:Odour Cue | 0.12 | 2 | .943 |

*Note.* p-values less than .05 shown in **bold**.

### 3.3 Sociability and Social Novelty Preference

Analysis of sociability in females from 3 – 12 months of age indicated that both transgenic females and wild-type controls preferred social over non-social interactions (Figure 3A and Table 5), but there were no effects of genotype, age, or a genotype by age interaction for preference ratio (Figure 3B and Table 5). For binned data at 12 months of age, there were no effects of genotype and no genotype by bin interaction for preference ratio (data not shown). For total investigation time, there was a genotype by age interaction for the sociability task (χ^2^ (1) = 4.91, *p* = .026), with transgenic females exhibiting reduced total investigation time (social and non-social investigation) compared to wild-type controls at 9 and 12 months of age (Cohen’s d 95% CIs [−0.46, 0.93], [−0.89, 0.50], [−1.40, 0.05], and [−1.65, −0.09] for genotype effects on investigation durations at 3, 6, 9, and 12 months of age, respectively) (Figures 3C, 3D, and 3E and Table 5). There was also a main effect of genotype (χ^2^ (1) = 7.68, *p* = .006) for time spent investigating both stimuli, with transgenic females exhibiting less investigation than wild-type controls. For binned data at 12 months of age, there was an overall effect of genotype for total investigation duration (χ^2^ (1) = 6.23, *p* = .013), but no genotype by bin interaction (Table 5). There was also no genotype effect or genotype by stimulus interaction for latency to first approach to the stimuli (data not shown).

**Figure 3.**
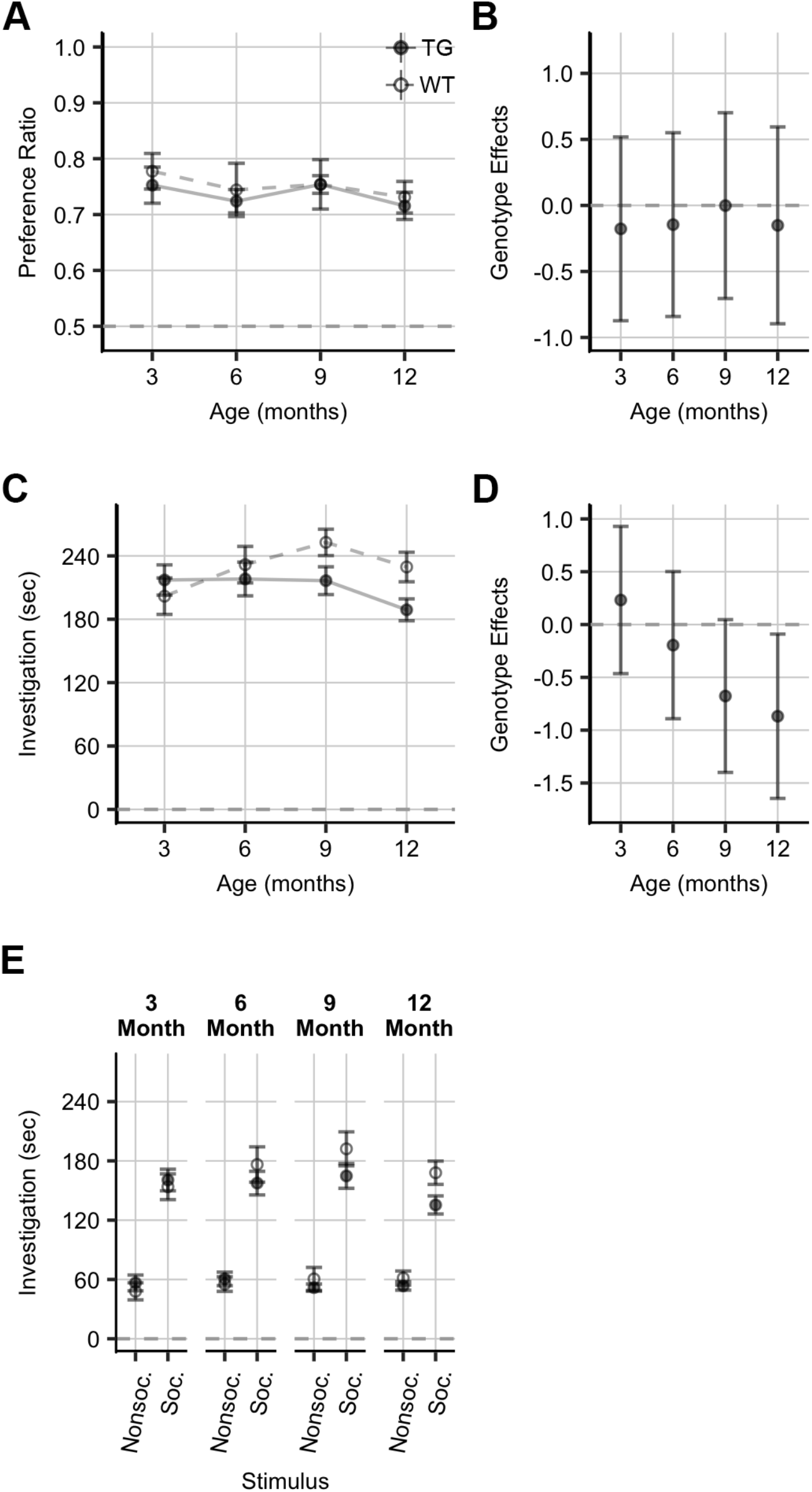
Overall Investigation in the Sociability Task. Mean and standard error for **A)** total investigation time (sec), **C)** preference ratio, and **E)** investigation time per side (sec), as well as 95% CIs for Cohen’s d for genotype effects for **B)** total investigation time (sec), and **D)** preference ratio, during the Sociability task for females from 3 to 12 months of age. Points indicate means, error bars indicate standard error (panels A, C, and E) and 95% CIs (panels B and D). Effects given for transgenic females relative to wild-type controls. TG = transgenic, WT = wild-type.

**Table 5.** Analysis of Deviance for the Sociability and Social Novelty Preference Tasks from 3 to 12 Months of Age

| Effect | Preference Ratio |  |  | Total Investigation Time |  |  |
| --- | --- | --- | --- | --- | --- | --- |
| | $\chi^2$ | df | *p* | $\chi^2$ | df | *p* |
| Sociability |  |  |  |  |  |  |
| Genotype | 0.27 | 1 | .601 | 7.68 | 1 | **.006** |
| Age | 1.01 | 1 | .316 | 0.01 | 1 | .913 |
| Genotype:Age | 0.06 | 1 | .805 | 4.91 | 1 | **.027** |
| Social Novelty Preference |  |  |  |  |  |  |
| Genotype | 2.68 | 1 | .102 | 0.05 | 1 | .819 |
| Age | 0.00 | 1 | .969 | 7.25 | 1 | **.007** |
| Genotype:Age | 0.04 | 1 | .846 | 2.54 | 1 | .111 |

Note. p-values less than .05 shown in **bold**.

Analysis of social novelty preference in females from 3 – 12 months of age indicated that both transgenic females and wild-type controls preferred the novel over familiar stimulus mouse (Figure 4A and Table 5), but there were no effects of genotype, age, or a genotype by age interaction for preference ratio (Figure 4B and Table 5). For binned data at 12 months of age, there were no effects of genotype and no genotype by bin interaction for preference ratio (data not shown). For total investigation time, there was a main effect of age (χ^2^ (1) = 7.25, *p* = .007), with both transgenic and wildtype females exhibiting increased investigation of both stimuli with age (95% CIs [143.85, 180.44], [177.02, 221.05], [187.79, 233.45], and [173.68, 215.88] for mean total investigation time at 3, 6, 9, and 12 months of age, respectively) (Figures 4C, 4D, and 4E and Table 5). There were no main effects of genotype, and no genotype by age interaction for total investigation time (Table 5). For binned data at 12 months of age, there were no effects of genotype and no genotype by bin interaction for total investigation time, and there were no genotype effects or genotype by stimulus interaction for latency to first approach to the stimuli (data not shown).

**Figure 4.**
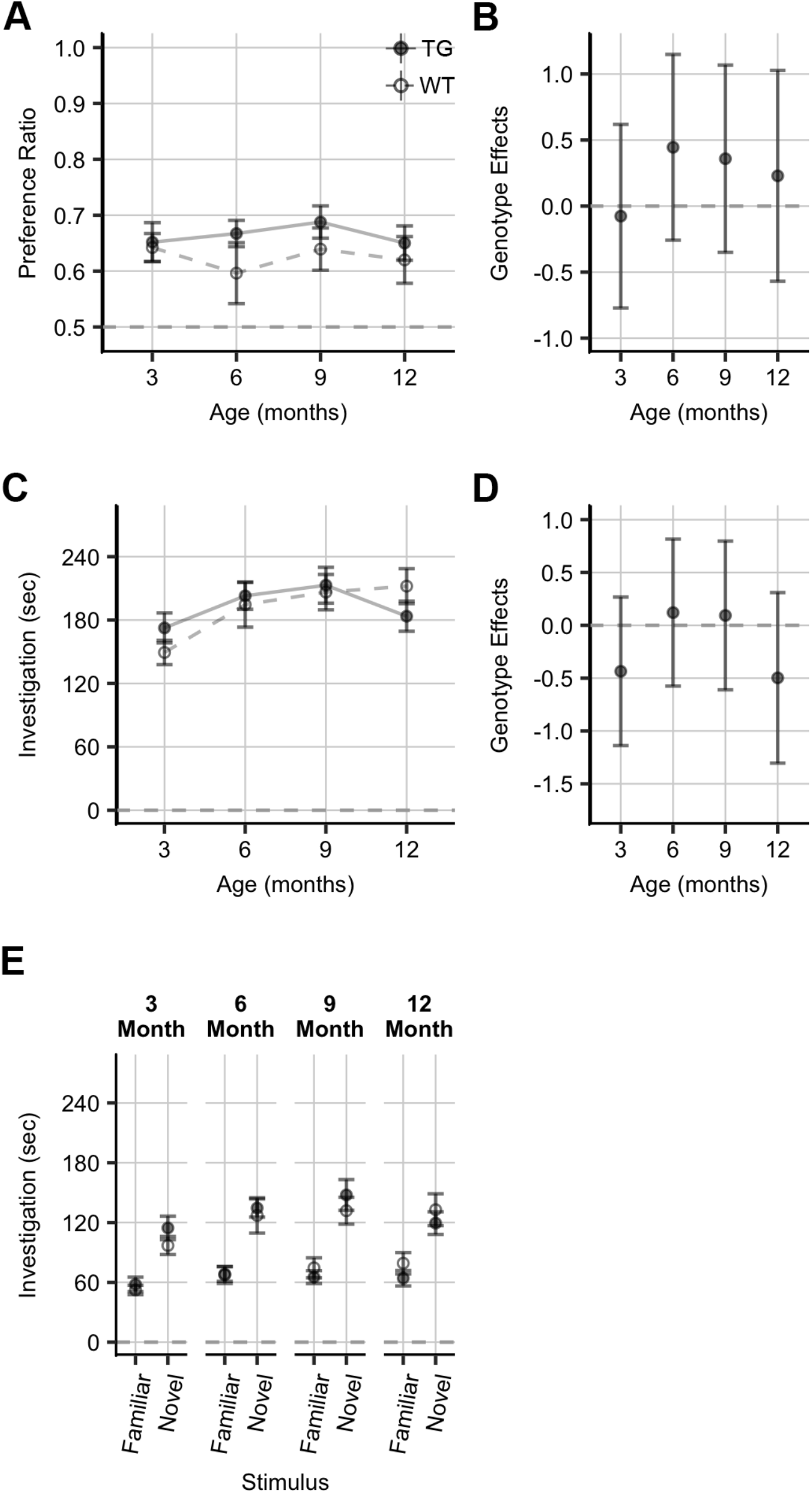
Social Investigation in the Social Novelty Preference Task. Mean and standard error for **A)** total investigation time (sec), **C)** preference ratio, and **E)** investigation time per side (sec), as well as 95% CIs for Cohen’s d for genotype effects for **B)** total investigation time (sec), and **D)** preference ratio, during the Social Novelty Preference task for females from 3 to 12 months of age. Points indicate means, error bars indicate standard error (panels A, C, and E) and 95% CIs (panels B and D). Effects given for transgenic females relative to wild-type controls. TG = transgenic, WT = wild-type.

### 3.4 Free Social Interaction

Analysis of behaviours in females from 3 – 12 months of age indicates a genotype by age interaction for orofacial sniffing (χ^2^ (1) = 4.14, *p* = .042), with transgenic females exhibiting decreased orofacial sniffing durations with age compared to wild-type controls (Cohen’s d 95% CI [−0.78, 0.56], [−1.21, 0.19], [−2.06, −0.51], and [−1.47, 0.08] for genotype effects at 3, 6, 9, and 12 months of age, respectively) (Figures 5A and 5B and Table 6). There was also a genotype by age interaction for following (χ^2^ (1) = 5.15, *p* = .023), with transgenic females exhibiting reduced following durations with age compared to wild-type controls (Cohen’s d 95% CIs [−0.55, 0.80], [−1.39, 0.02], [−1.51, −0.04], and [−1.85, −0.24] for genotype effects at 3, 6, 9, and 12 months of age, respectively) (Figures 5C and 5D and Table 6). There was a main effect of genotype on anogenital sniffing (χ^2^ (1) = 4.52, *p* = .034); Cohen’s d 95% CI [−0.79, −0.10]), with transgenic females exhibiting decreased durations compared to wild-type controls (Figures 5E and 5F and Table 6). For binned data at 12 months of age, there was a genotype by bin interaction for orofacial sniffing (χ^2^ (1) = 7.46, *p* = .006), anogenital sniffing (χ^2^ (1) = 4.35, *p* = .039), and following behaviour (χ^2^ (1) = 8.86, *p* = .003), with transgenic females exhibiting reduced durations of each behaviour during the first two minutes of the trial (Table 6). There were no effects for self-grooming behaviour (Figure 5G and 5H and Table 6).

**Figure 5.**
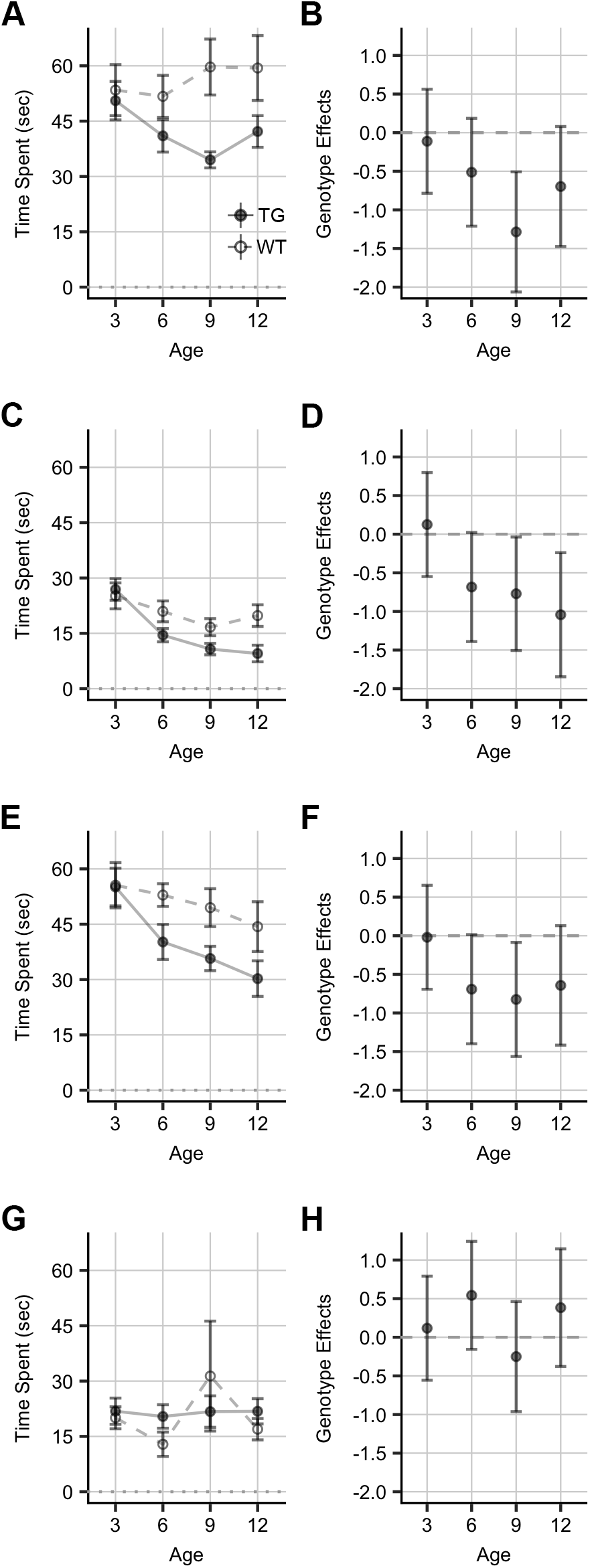
Social Investigation in the Free Social Interaction Task. Mean and standard error for duration (in sec) of **A)** orofacial sniffing, **C)** following, **E)** anogenital sniffing, and **G)** self-grooming behaviours, as well as 95% CIs for Cohen’s d for genotype effects for **B)** orofacial sniffing, **D)** following, **F)** anogenital sniffing, and **H)** selfgrooming behaviours, during the Free Social Interaction task for females from 3 to 12 months of age. Points indicate means, error bars indicate standard error (panels A, C, E, and G) and 95% CIs (panels B, D, F, and H). Effects given for transgenic females relative to wild-type controls. TG = transgenic, WT = wild-type.

**Table 6.** Analysis of Deviance for the Free Social Interaction Task from 3 to 12 Months of Age

| Effect | $\chi^2$ | df | *p* |
| --- | --- | --- | --- |
| Orofacial Sniffing |  |  |  |
| Genotype | 0.69 | 1 | .406 |
| Age | 31.85 | 1 | **.000** |
| Genotype:Age | 7.46 | 1 | **.006** |
| Anogenital Sniffing |  |  |  |
| Genotype | 0.64 | 1 | .422 |
| Age | 75.13 | 1 | **.000** |
| Genotype:Age | 4.25 | 1 | **.039** |
| Follow |  |  |  |
| Genotype | 1.90 | 1 | .168 |
| Age | 53.52 | 1 | **.000** |
| Genotype:Age | 8.86 | 1 | **.003** |
| Self-Grooming |  |  |  |
| Genotype | 0.48 | 1 | .487 |
| Age | 2.95 | 1 | .086 |
| Genotype:Age | 0.19 | 1 | .661 |

*Note.* *p*-values less than .05 shown in **bold**.

## 4. Discussion

Compared to wild-type controls, transgenic 5xFAD females exhibited reduced interest in social odours during the olfactory habituation/dishabituation task, but no obvious olfactory deficits and no changes in scent marking behaviour compared to wild-type controls. Transgenic females also displayed an age-related decrease in investigation in the sociability and free social interaction tasks, although no differences during the social novelty preference task were observed at any age. This suggests that transgenic 5xFAD females exhibit impaired social interaction with age, but that this change is context-specific.

### 4.1 Transgenic 5xFAD Mice Exhibit Decreased Interest in Social Odours, but Normal Olfactory Perception and Scent-based Social Communication

Transgenic 5xFAD females from 3 – 12 months of age showed no observable signs of non-social olfactory deficits relative to wild-type controls. The overall pattern of responses on the olfactory habituation/dishabituation task indicates that transgenic mice are able to detect non-social odours and discriminate between two different non-social odours (Figures 1 and S1). Previous findings on olfactory perception in 5xFAD mice have been mixed; prior reports support our findings that female 5xFAD mice have no overall deficits in olfactory perception [38-40], although one study has reported that transgenic 5xFAD male mice display age-related olfactory deficits beginning at 3 months of age [41].

Transgenic 5xFAD females exhibited decreased interest in social odour cues by 3 months of age, and this persisted through to 12 months of age (Figure 1). Previous research indicates that females in the 3xTg-AD mouse strain exhibit decreased preference for male social odours compared to wild-type controls, although these mice also exhibited a concomitant impairment in odour discrimination [42]. This suggests that the olfactory deficits for male social odours demonstrated by female transgenic 3xTG-AD mice may be the result of overall deficits in olfaction, while the social olfactory deficit exhibited here by transgenic 5xFAD females are due to reduced motivation to explore social odours specifically.

While there was minimal dishabituation to the second social odour during the olfactory habituation/dishabituation task, the same effect was seen for both transgenic and wild-type females. This suggests that all experimental mice had difficulty discriminating between odour cues from same-sex mice from the same strain, most likely due to similarities between the chemical characteristics of the odours (e.g., major histocompatibility complex and/or major urinary proteins) [28,43,44]. These proteins are associated with determination of individuality in animals and are influenced by both genetics and environment, and thus, they are likely to be very similarly expressed in our stimuli odours, which were all taken from cages housing highly inbred 5xFAD mice who have been housed in nearly identical conditions.

Although transgenic mice were hypothesized to exhibit impaired responses to social odour cues, the pattern of urine marking in response to social vs non-social odour cues was comparable between transgenic females and wild-type controls at all ages (Figure 2). Additionally, the overall increase in urine marking in response to social compared to non-social odour cues indicates that transgenic mice are able to discriminate between these cues and mediate their response accordingly, thereby indicating no deficits in this form of scent-based communication. While these normal levels of urine marking may seem counter to the decreased interest in social odour cues exhibited during the olfactory habituation/dishabituation task, they are similar to effects seen in the transgenic BTBR model of autism spectrum disorder. Compared to wild-type C57Bl/6 controls, BTBR males and females at ~3 months of age exhibit reduced sociability [45] and an overall decrease in sniffing durations during the olfactory habituation/dishabituation task [35], while exhibiting similar levels of countermarking in response to urine from novel male conspecifics [26]. While these results are not directly comparable to mouse models of AD, they do indicate that interest in social odours can be dissociated from social communication via urine marking, and that this form of social communication is distinct from sociability. Our findings indicate that transgenic 5xFAD females exhibit decreased motivation for investigating social odours but are able to maintain normal urine marking behaviours.

### 4.2 Transgenic 5xFAD Females Exhibit Reduced Social Approach During Novel Situations

Transgenic 5xFAD females exhibited an age-dependent decrease in total investigation time (social and non-social) compared to wild-type controls during the sociability trial in the three-chamber apparatus (Figures 3C and 3D), but there were no genotype effects during the subsequent social novelty preference trial (Figures 4C and 4D), suggesting that this effect is specific to novel situations. Transgenic females and wild-type controls exhibit similar preference ratios for social over non-social interactions (Figures 3A and 3B), and novel over familiar interactions (Figures 4A and 4B), indicating that transgenic females are still interested in social interactions in this context. However, transgenic 5xFAD females spent less time engaging in social investigative behaviours during the free social interaction task (Figure 5). Together, these findings suggest that transgenic 5xFAD females exhibit increased social anxiety compared to wild-type controls when exposed to reciprocal social contact. The initial exposure to a novel conspecific during the sociability and free social interaction tasks could result in a high level of social anxiety, leading to reduced investigative behaviours overall.

Although transgenic 5xFAD mice do not exhibit increased general anxiety compared to wild-type controls [24,38,46,47], evidence from the Fmr1-KO strain (a model of Fragile X Syndrome) suggests general anxiety can be dissociated from social anxiety. Fmr1-KO mice exhibit decreased anxiety in open field and light/dark tests, yet increased social anxiety on the mirrored-chamber test, a test that uses a mirrored chamber to create the visual illusion that another mouse is present, and compares exploration to this chamber to chambers without mirrors [48]. Spencer *et al* [48] also indicate that treatment with anxiolytics reduces the latency of mice to enter the mirrored chamber, suggesting its validity as a method for assessing social anxiety. Fmr1-KO mice also exhibit a preference for social over non-social interactions, with a decrease in social interactions during the second half of a free social interaction task [49]. Although the transgenic 5xFAD females in the present study exhibited decreased investigation durations compared to wild-type controls at the beginning of the trial, rather than the end, the results obtained with Fmr1-KO mice indicate that transgenic mice may be interested in social interaction but less willing than wild-type controls to approach free-roaming novel conspecifics, and that these effects may arise due to different levels of social—but not general—anxiety.

Overall, this pattern of behaviour—consistent social approach to a restricted mouse but an age-related decrease in investigative behaviour during reciprocal interactions—has not been reported in other mouse models of AD. Transgenic APP/PS1 males at 6 months of age exhibit similar levels of social approach behaviour as wild-type controls in the three-chamber apparatus, but high levels of non-social exploration [18]. Compared to wild-type controls, transgenic 3xTg-AD females exhibit fewer social behaviours during a free social interaction task at 18 months of age, a deficit that was preceded by higher levels of social interaction during free social interaction at 12 months of age [19]. Transgenic Tg2576 females at 21 months of age exhibited no difference in social investigation compared to wildtype controls during free social interaction [17]. Thus, social behaviours in mouse models of AD appear to be strain dependent, and previous research has not performed both the three-chamber sociability test and the free social interaction test in the same animals, making it difficult to assess whether the particular combination of effects observed in the current study is unique to the 5xFAD strain.

## 5. Conclusions

The 5xFAD mouse model provides a unique opportunity to study the effects of early-, mid- and late-stages of brain pathology related to Alzheimer’s disease. While a recent review suggested that APP-overproducing models typically do not exhibit progressive age-related behavioural deficits [50], our longitudinal study demonstrates that the limited deficits in social-related behaviours demonstrated by transgenic 5xFAD females at 3 months of age do progress as they age. Transgenic 5xFAD female mice show an age-related decrease in investigation times in the sociability and free social interaction tasks, as well as a persistent decrease in social interest during the olfactory habituation/dishabituation task. Transgenic 5xFAD females exhibit no overall changes in scent marking behaviour relative to wild-type controls at any age, as well as no differences in preference ratios during the sociability or social novelty preference tasks in the three-chamber apparatus, suggesting that social deficits in 5xFAD transgenic female mice are most evident during high-arousal situations, and may be mitigated by habituation to an environment. Overall, the progressive change in social investigation exhibited by transgenic 5xFAD females mimics some aspects of the social withdrawal observed in human AD patients, suggesting that this strain may be suitable for modelling the social dysfunction observed in human patients over disease progression; however, further exploration is needed to better define how context contributes to changes in social function observed in 5xFAD mice.

## 6. Acknowledgements

The authors would like to thank Dr. Richard Brown for his valuable input on the described experiments, and Lindsey Colyn for her help with behavioural testing.

## 7. Funding Sources

This work was supported by the Alzheimer’s Association [grant number 2016-MNIRGD-391961] and the Canadian Foundation for Innovation and Nova Scotia Research and Innovation Trust [grant number 34123].

## 8. Declarations of Interest

None.

## Supplementary Figure Caption

**Figure S1.**
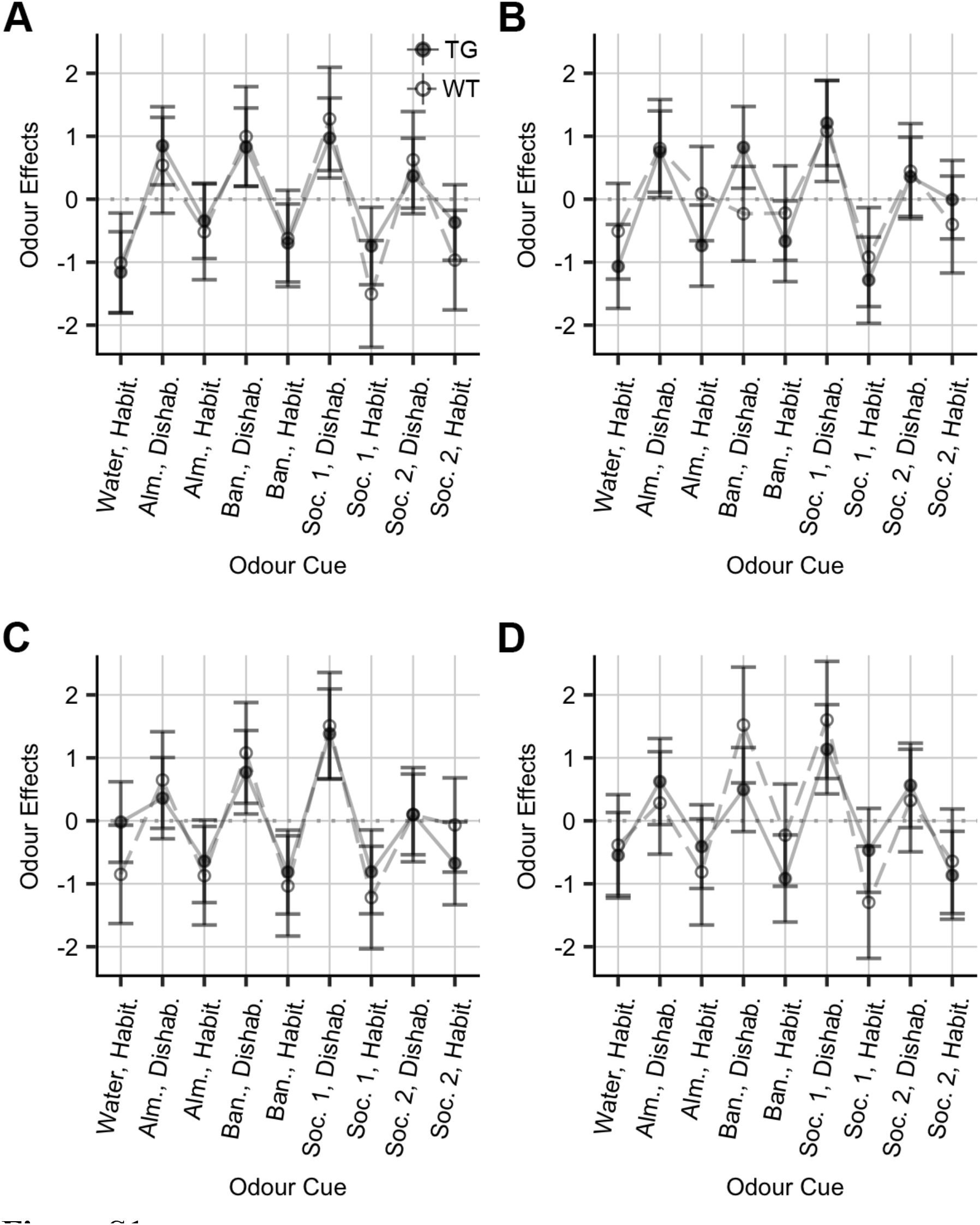
Odour and Presentation Effects in the Olfactory Habituation/Dishabituation Task. 95% CIs for Cohen’s d for habituation (Habit.) and dishabituation (Dishab.) effects for each odour during the Olfactory Habituation/Dishabituation task for females at **A)** 3 months, **B)** 6 months, **C)** 9 months, and **D)** 12 months of age. Points indicate means, error bars indicate 95% CIs. TG = transgenic, WT = wild-type, Alm. = almond cue, Ban. = banana cue, Soc. 1 = first social odour cue, Soc. 2 = second social odour cue.

